# Self-antigen driven affinity maturation is required for pathogenic monovalent IgG4 autoantibody development

**DOI:** 10.1101/2020.03.14.988758

**Authors:** Miriam L. Fichtner, Casey Vieni, Rachel L. Redler, Ljuvica Kolich, Ruoyi Jiang, Kazushiro Takata, Panos Stathopoulos, Pablo Suarez, Richard J. Nowak, Steven J. Burden, Damian C. Ekiert, Kevin C. O’Connor

## Abstract

Pathogenic IgG4 autoantibodies in autoimmune myasthenia gravis (MG) are functionally monovalent as a result of Fab-arm exchange. The origin and development of these unique autoantibodies are not well understood. We examined MG patient-derived monoclonal autoantibodies (mAbs), their corresponding germline-encoded unmutated common ancestors (UCA) and monovalent antigen-binding fragments (Fabs) to investigate how antigen-driven affinity maturation contributes to both binding and immunopathology. Mature mAbs, their UCA counterparts and mature monovalent Fabs bound to the autoantigen and retained their pathogenic capacity. However, monovalent UCA Fabs still bound the autoantigen but lost their pathogenic capacity. The mature Fabs were characterized by very high affinity (sub-nanomolar) driven by a rapid on-rate and slow off-rate. However, the UCA affinity was approximately 100-fold less than that of the mature Fabs, which was driven by a rapid off-rate. Crystal structures of two Fabs shed light on how mutations acquired during affinity maturation may contribute to increased MuSK binding affinity. These collective findings indicate that the autoantigen initiates the autoimmune response in MuSK MG and drives autoimmunity through the accumulation of somatic hypermutation such that monovalent IgG4 Fab-arm exchanged MG autoantibodies reach a high affinity threshold required for pathogenic capacity.

**Summary:** IgG4 autoantibodies in autoimmune myasthenia gravis are functionally monovalent, requiring a high affinity threshold featuring fast on/slow off rates, to reach pathogenic capacity. This capacity is dependent on self-antigen initiated and driven maturation, which includes the accumulation of indispensable somatic hypermutations that may alter electrostatic interactions with the antigen.

## Introduction

Myasthenia gravis (MG) is a chronic autoimmune disorder affecting neuromuscular transmission [1, 2]. The disease is caused by pathogenic autoantibodies that target components of the neuromuscular junction. Given that the immunopathogenesis is directly governed by known autoantibody-autoantigen combinations, MG serves as an archetype for B cell-mediated autoimmune disease. MG disease subsets are classified by autoantibody specificity; autoantibodies to the acetylcholine receptor (AChR) [3] are found in most patients, followed by autoantibodies to muscle-specific tyrosine kinase (MuSK) in other patients [4]. The clinical presentation among the subtypes is often similar, but the underlying immunopathology is decidedly divergent. The MuSK subtype highlights this distinction as the autoantibodies in MuSK MG are primarily IgG4 [5], a subclass that does not share key properties found in the other subclasses. The most intriguing feature of human IgG4 antibodies is their unique ability to participate in “Fab-arm exchange”, such that a monospecific IgG4 antibody exchanges a heavy and light chain pair with another IgG4 antibody to become bispecific [6]. Consequently, IgG4 antibodies are asymmetric antibodies with two different antigen-combining sites and therefore possess monovalent specificities. Serum IgG4 autoantibodies that have undergone Fab-arm exchange contribute to the pathology of MuSK MG [7].

During the course of a developing immune response to an exogenous antigen, B cells produce antibodies with increased affinity as they proceed through the process of affinity maturation [8–10]. The successively greater antibody affinities accumulate as a direct result of clonal selection and somatic hypermutation (SHM). B cell responses to self-antigen in most human autoimmune diseases appear to be products of this affinity maturation process. Autoantibodies with pathogenic capacity, isolated from patients with neuromyelitis optica (NMO), pemphigus vulgaris (PV) or AChR MG are characterized by the hallmarks of this process including the accumulation of somatic hypermutation [11–13]. Recently, cloned autoantibodies that target MuSK were isolated from patients with MG [14–16]. These autoantibodies include the hallmarks of affinity maturation, including a high frequency of somatic hypermutation. Given that IgG4 antibodies are often the product of a response to chronic exposure to exogenous antigens [17], such as allergens, it is not clear whether these autoantibodies are produced by B cells that were driven through the affinity maturation process by the autoantigen, MuSK. Moreover, MuSK MG autoantibodies are functionally monovalent, as a consequence of Fab-arm exchange, thus the binding does not benefit from the accumulated strength of multiple affinities (avidity) that bivalent antibodies use to their advantage. Thus, the affinity threshold for functional binding and pathogenic capacity may be higher than that of other autoantibodies and may consequently be highly dependent on affinity maturation.

We sought to further understand whether a self-antigen was driving the autoimmune response in MuSK MG. In addition, we determined how somatic hypermutation contributes to MuSK autoantibody binding and pathogenic capacity in the context of both divalent and monovalent binding. We carried-out these experiments by examining a set of MuSK MG-derived human recombinant monoclonal antibodies (mAbs). These mAbs were reverted to their unmutated common ancestors (UCAs) by replacing all of the acquired somatic mutations with germline encoded residues. They were expressed and evaluated as divalent mAbs or monovalent antigen-binding fragments (Fabs), the latter of which allowed us to directly test the properties of Fab-arm exchanged products. We found that both mature and germline encoded mAbs bound to MuSK and retained their pathogenic capacity. The mature monovalent Fabs also bound MuSK and demonstrated pathogenic capacity. However, the germline encoded Fabs bound MuSK but lost their pathogenic capacity. Affinity measurements revealed that the mature Fabs possessed very high affinity for MuSK, driven by a rapid on-rate and slow off-rate. The affinities of the germline encoded Fabs were approximately 100-fold less than the mature Fabs, largely due to a rapid off-rate. Crystal structures of two Fabs revealed that acquired mutations in the mature Fabs increase the negative charge of the antigen binding region, likely contributing to increased binding affinity to potential positively charged epitopes on MuSK. Taken together, these findings indicate that the autoantigen, MuSK, both initiates and drives the autoimmune response in MuSK MG. They furthermore demonstrate that IgG4 Fab-arm exchanged MuSK autoantibodies require high affinity binding to reach pathogenic capacity, demonstrating a particularly critical role for somatic hypermutation in these unique monovalent autoantibodies.

## Results

### Unmutated common ancestors (UCA) of MuSK mAbs bind to the autoantigen MuSK

We previously isolated single B cells that expressed MuSK autoantibodies from two MuSK MG patients. From these single B cells, we produced human recombinant monoclonal autoantibodies (mAb) (MuSK1A, 1B and 3-28) that bound to MuSK and demonstrated pathogenic capacity **(Supplemental Table 1)** [15, 16]. Here, we sought to investigate how affinity maturation contributed to MuSK binding. To this end, we reverted the mAbs to their germline configuration. Amino acid residue changes, arising from the somatic hypermutation (SHM) process, in the heavy and light variable region gene segments, including the CDR3s, were identified and mutated back to the unmutated common ancestor (UCA) sequence configuration using a step-wise approach **(Supplemental Figure 1)**. This procedure generated a series of mAbs in which only individual regions (CDRs and FRs), or combinations of regions were changed to the germline configuration. This series of partially reverted mAbs (intermediate reversions) provided the opportunity for us to evaluate how SHM affected binding in specific regions. The mature mAbs, the intermediate reversions and the UCAs were tested for their ability to bind MuSK using a live cell-based assay (CBA).

We compared the mature mAbs to their complete UCA counterparts **(Figure 1 A and B)**. These mAbs, as well as negative and positive control mAbs, were tested over a range of concentrations (10 – 0.02 µg/mL) **(Figure 1 B)**. The mean fluorescent intensity (MFI) of MuSK antigen transfected cells was subtracted from the MFI of non-transfected cells (ΔMFI). The UCAs of MuSK1A and MuSK1B antibodies and their mature counterparts bound MuSK similarly: all four mAbs showing positive binding at concentrations as low as 0.02 µg/mL. The UCA of autoantibody MuSK3-28 showed diminished binding over this concentration range, compared to its mature counterpart and was positive only at 10 – 0.2 µg/mL, while the mature mAb remained positive at 10 – 0.02 µg/mL.

**Figure 1.**
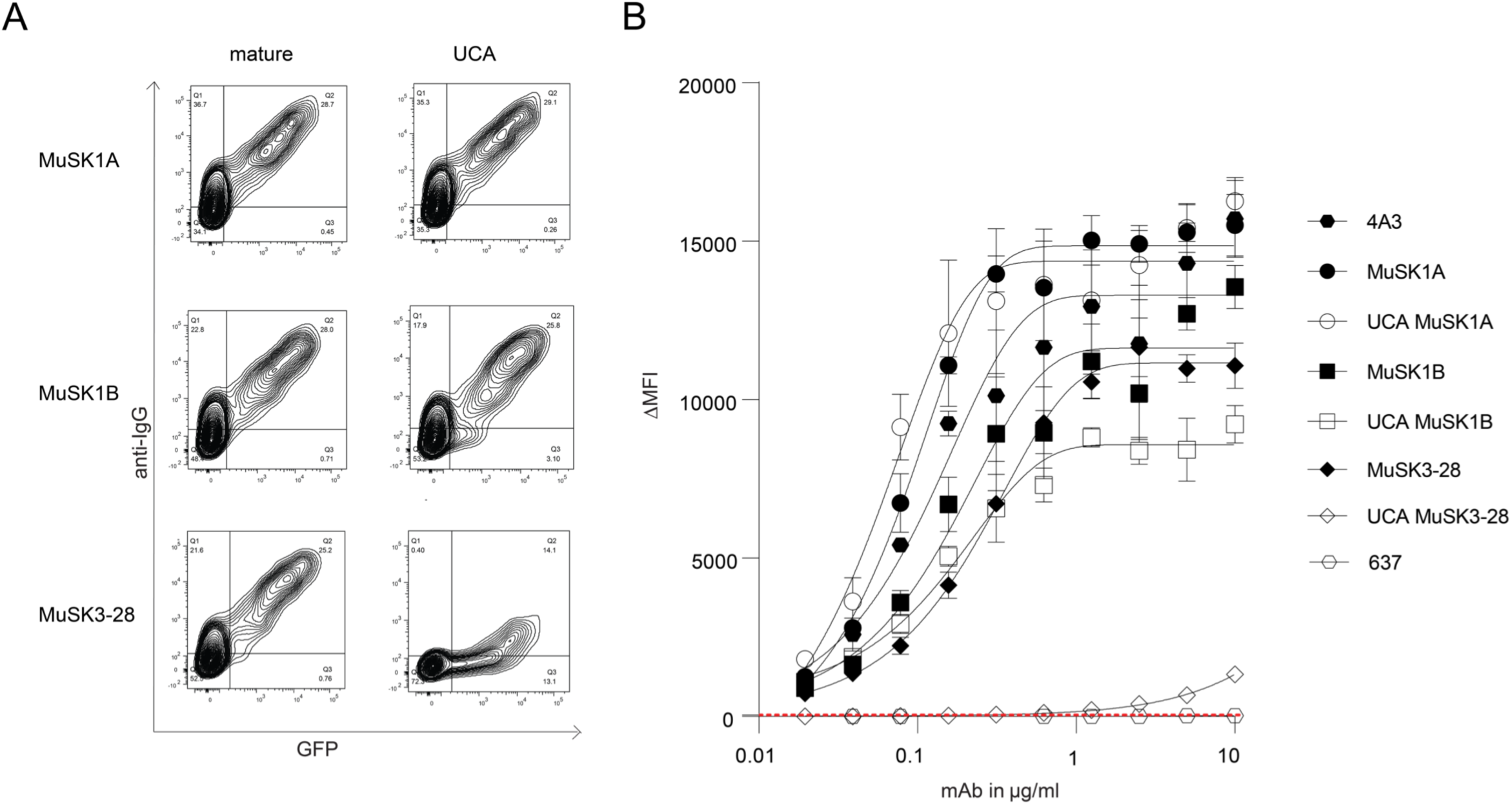
Unmutated common ancestors of MuSK mAbs bind to the MuSK autoantigen. MuSK-specific mAbs and their UCAs were tested for surface binding to MuSK on MuSK-GFP– transfected HEK cells. **(A)** Representative cell-based assay (CBA) contour plots are shown for the three MuSK-mAbs and their UCA. The x-axis represents GFP fluorescence intensity and, consequently, the fraction of HEK cells transfected with MuSK. The y-axis represents Alexa Fluor 647 fluorescence intensity, which corresponds to secondary anti–human IgG Fc antibody binding and, consequently, primary antibody binding to MuSK. Hence, transfected cells are located in the right quadrants and cells with MuSK antibody binding in the upper quadrants. The plots show testing with a mAb concentration of 1.25 µg/ml. **(B)** Binding to MuSK was tested over a series of ten two-fold dilutions of each mAb ranging from 10-0.02 µg/ml. Humanized MuSK mAb 4A3 was used as the positive control and AChR-specific mAb 637 as the negative control. The ΔMFI was calculated by subtracting the signal from non-transfected cells from the signal of transfected cells. Each data point represents a separate replicate within the same experiment, which was performed in triplicate. Bars or symbols represent means and error bars SDs. Values greater than the mean + 4SD of the negative control mAb at 1.25 µg/ml (indicated by the horizontal dotted line) were considered positive.

Given that the UCA of mAb MuSK3-28 demonstrated diminished binding to MuSK, we next explored the binding of the intermediate reversion mAbs to evaluate how SHMs in the individual CDRs and FRs contributed to binding. We tested the intermediate reversions mAbs of MuSK1A, MuSK1B and MuSK3-28, together with both negative and positive control mAbs over a range of mAb concentrations (10 – 0.02 µg/mL). Consistent with the UCA binding, all intermediate constructs tested for MuSK1A bound MuSK similar to the mature mAb **(Supplemental Figure 2 A)**. A number of the intermediates for MuSK1B showed a minor decrease in binding to MuSK. However, the binding remained 1639 to 2183-fold above background at the concentration of 1.25 µg/mL, which we previously used to distinguish between binders and non-binders [16] (**Supplemental Figure 2 B**). Conversely, intermediate constructs for mAb MuSK3-28 showed more pronounced changes in binding. The largest cause for diminished binding of MuSK3-28 could be attributed to a reversion of the H CDR1 and H CDR2 domain together with the H FR1, located close to the H CDR1 region of the heavy chain. Other reversions with either H CDR1 and H CDR2 alone or in combination with the H FR2 region did not show a similar impact on binding. The ΔMFI of the mature mAb was 7824, while the HCDR12FR1 reversion was 75 (104-fold change) at 1.25 µg/mL **(Supplemental Figure 3 A)**.

We included a non-MuSK MG mAb to serve as a control. We reverted mAb 637 (a MG patient-derived recombinant monoclonal that recognizes AChR, [12]) back to the germline encoded UCA sequence. A CBA specific for AChR was used to assess the binding to the AChR. The heavy chain CDR2 region germline reversion of mAb 637 showed diminished binding compared to the mature mAb 637 **(Supplemental Figure 3 B)**. These results were reproducible over a broad range of concentrations. In summary, these findings demonstrate that UCA counterparts of the three MuSK mAbs all bind to MuSK.

### The UCA bind to the same MuSK domain, are not polyspecific and the autoantibody light chains contribute to MuSK binding

The three mature mAbs MuSK1A, MuSK1B and MuSK3-28 recognized an epitope present in the Ig2-like domain of MuSK [16]. We tested whether the UCA antibodies recognize the same epitope as their mature counterparts, using several variants of the MuSK antigen. The extracellular region of MuSK is comprised of three Ig-like domains and a frizzled-like domain **(Supplemental Figure 4 A)**. Both the mature and UCA mAbs recognized the same domain (Ig2-like) on MuSK **(Supplemental Figure 4 B and C)**.

We next tested whether binding of the UCA mAbs was attributable to polyspecificity. Using a well-established approach, we tested reactivity against LPS, dsDNA and insulin by ELISA, as binding to all three of these antigens is a property of polyspecific antibodies [18]. Binding to all three antigens was not observed for the three mature MuSK mAbs or their UCA counterparts **(Supplemental Figure 5)**.

We subsequently investigated whether the MuSK mAb light chains contribute to binding, or if the binding is heavy chain dependent only. Therefore, all three mature MuSK mAb heavy chains were paired with non-endogenous light chains that were not originally paired with the heavy chain and tested for binding to MuSK by CBA. All of the mAbs with light chain swaps showed significantly diminished binding (p-value range 0.01-0.0001) compared to the endogenous pair **(Supplemental Figure 6 A-C)**. In summary, both the mature and UCA mAbs recognize the same domain (Ig2-like) on MuSK, lack polyspecificity and the endogenous pairing of the heavy and light chain is important for binding.

### The UCA mAbs have lower affinity for MuSK than their mature counterparts

We next sought to measure the affinities of the antibodies to quantify the strength of the interaction between the MuSK mAbs and UCA. We produced monovalent Fabs to measure affinity, rather than avidity, and also to emulate the functional monovalency of Fab-arm exchanged IgG4. All three mature Fabs displayed high affinity for MuSK. Fab MuSK1A (K_D_ = 0.41 nM) and Fab MuSK1B (K_D_ = 0.44 nM) bound to MuSK with 30 times higher affinity than Fab MuSK3-28 (K_D_ = 12 nM) **(Supplemental Figure 7 A, C and E)**. The UCAs of all three mature Fabs had a lower affinity for MuSK than the mature Fabs **(Table 1)**. The affinity of UCA Fab MuSK1A for MuSK dropped 76-fold (K_D_ = 31 nM); the affinity of UCA Fab MuSK1B dropped 120-fold (K_D_ = 53 nM); the affinity of UCA Fab MuSK3-28 decreased by 73-fold (K_D_ = 870 nM) **(Supplemental Figure 7 B, D and F)**. The dissociation rates (k_off_) were remarkably faster for the UCA Fabs compared to their mature counterparts **(Table 2)**. The comparison of the UCA and mature Fab of MuSK3-28 **(Table 2)** showed a slower association rate and similar dissociation rate. In summary, the binding kinetics of the UCA and mature Fabs show altered dissociation rates that contribute to an approximate 100-fold decrease in affinity.

**Table 1.** K_D_ values of mature and UCA MuSK1A, MuSK1B and MuSK3-28.

|  | **K<sub>D</sub> of mature Fab (nM)** | **K<sub>D</sub> of unmutated ancestor Fab (nM)** | **Fold change** |
| --- | --- | --- | --- |
| **MuSK1A** | 0.41 ± 0.0045 | 31 ± 0.21 | 76 |
| **MuSK1B** | 0.44 ± 0.0040 | 53 ± 0.59 | 120 |
| **MuSK3-28** | 12 ± 0.022 | 870 ± 12 | 73 |

**Table 2.** k_on_ and k_off_ values of mature and UCA MuSK1A, MuSK1B and MuSK3-28.

| | $k_{\text{on}} (\text{M}^{-1} \text{s}^{-1})$ | $k_{\text{off}} (\text{s}^{-1})$ |
| --- | --- | --- |
| **MuSK1A** | $3.2 \pm 0.03 \times 10^5$ | $1.3 \pm 0.007 \times 10^{-4}$ |
| **UCA MuSK1A** | $5.7 \pm 0.04 \times 10^5$ | $1.8 \pm 0.003 \times 10^{-2}$ |
| **MuSK1B** | $4.3 \pm 0.04 \times 10^5$ | $1.9 \pm 0.008 \times 10^{-4}$ |
| **UCA MuSK1B** | $4.0 \pm 0.04 \times 10^5$ | $2.1 \pm 0.006 \times 10^{-2}$ |
| **MuSK3-28** | $1.4 \pm 0.003 \times 10^5$ | $1.6 \pm 0.001 \times 10^{-3}$ |
| **UCA MuSK3-28** | $0.022 \pm 0.0003 \times 10^5$ | $1.9 \pm 0.009 \times 10^{-3}$ |

### Crystal structures of MuSK1A and MuSK1B Fabs

To understand the structural basis for how affinity maturation contributes to MuSK binding we obtained crystals of MuSK1A and MuSK1B to 1.8 Å and 1.75 Å respectively **(Figure 2 A and B; Supplemental Table 2)**. The overall structure of both MuSK1A and MuSK1B were comparable to a number of Fabs with high CDR amino acid sequence similarity found in the Protein Data Bank (PDB) **(Supplemental Figure 8)**. Compared to these sequence-related Fabs, MuSK1A and MuSK1B differed most significantly in the heavy and light CDR3 loops, consistent with these loops being the sites of greatest diversification in most antibodies. For both MuSK1A and MuSK1B, the mutations away from the UCA sequence were distributed throughout the Fab variable domains with approximately 40-50% of the mutations clustered in the CDR loops, and the remainder scattered throughout the framework regions **(Figure 2 C and D)**. In the crystal structure for MuSK1B we observed blurred electron density in the vicinity of the heavy chain “elbow” region, which connects the variable (VH) and constant domain (CH1) of an antibody. Blurred electron density was observed for the linker between the VH and CH1 domains, as well as several nearby turns between strands of the VH domain regions. The lower quality of electron density in this local area likely indicated flexibility or multiple conformations within this region.

**Figure 2.**
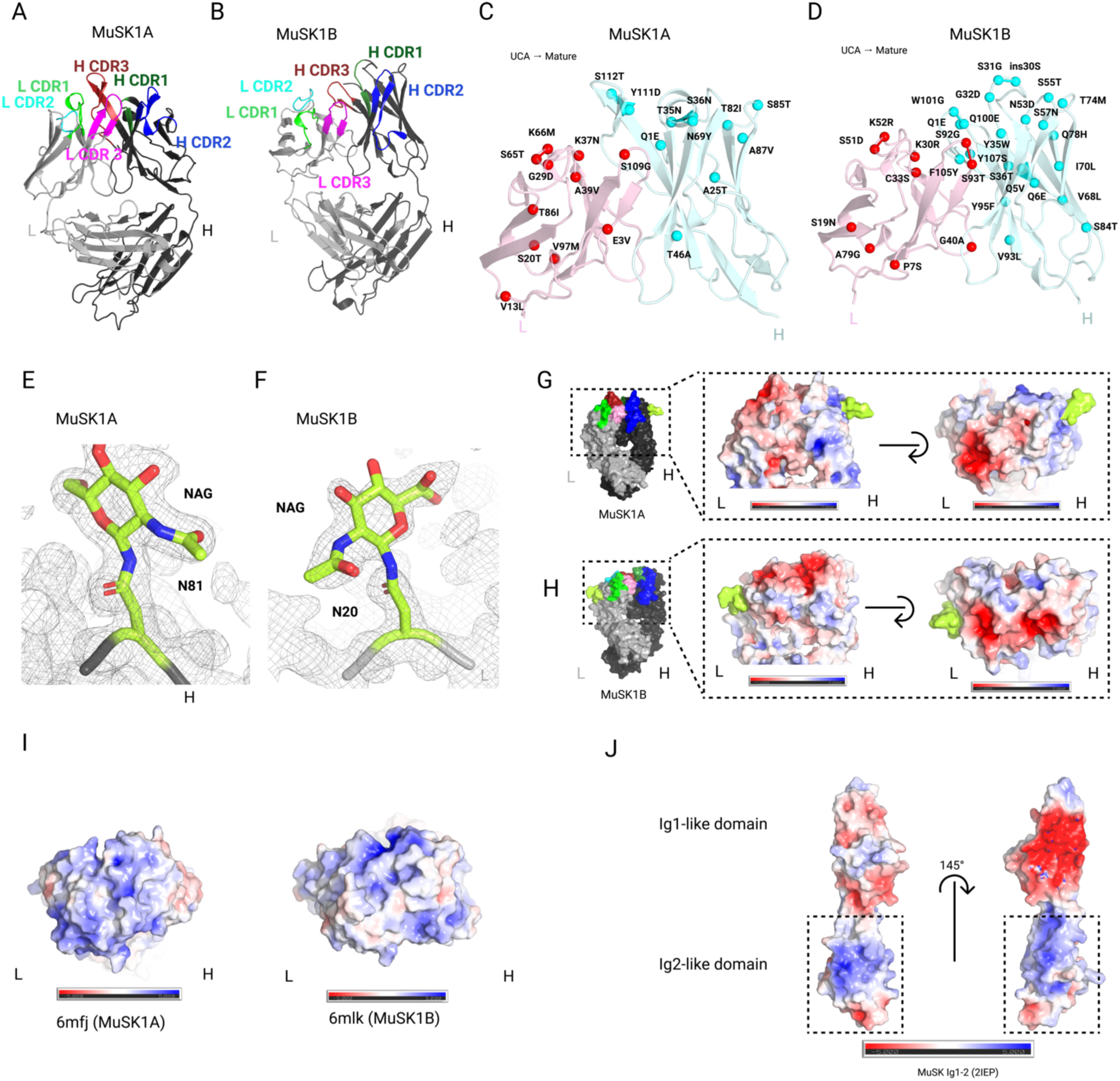
Crystal Structure, mutation map, glycosylation and electrostatic potential maps of MuSK1A and MuSK1B and MuSK Ig1-2 like domain. **(A)** Structure of MuSK1A heavy and light chains (*PDB code pending*). **(B)** Structure of MuSK1B heavy and light chains (*PDB code pending*). **(C)** UCA mutations mapped to the mature MuSK1A. UCA mutations are scattered throughout the mature MuSK1A CDR and FR regions. MuSK1A light chain is colored light pink and MuSK1A heavy chain is colored pale cyan. UCA mutants’ C_alpha_ carbons are shown as darker red or cyan spheres for the light or heavy chain respectively with the mutation from the UCA to the mature mAb. **(D)** UCA mutations mapped to mature MuSK1B. UCA mutations are uniformly distributed throughout the mature MuSK1B CDR loops and FR regions. **(E)** N81 glycosylation site in MuSK1A heavy chain at a threshold of 1.0 σ in the 2Fo-Fc map. MuSK1B heavy chain backbone is shown as a black tube. **(F)** N20 glycosylation site in the MuSK1B light chain at a threshold of 1.0 σ in the 2Fo-Fc map. MuSK1B light chain backbone is shown as a light gray tube. **(G and H)** Electrostatic potential map of MuSK1A and MuSK1B at the antigen binding site with the modeled glycans shown in yellow green. In MuSK1A the light chain is predominantly negatively charged, while in MuSK1B both the heavy and light chains are predominantly negatively charged. **(I)** Electrostatic potential map of sequence-related Fabs for UCA MuSK1A (6mfj) and UCA MuSK1B (6mlk) at the antigen binding site. **(J)** Electrostatic potential map of the MuSK Ig1-like and Ig2-like domain. The MuSK Ig1-like domain is predominantly negative, while MuSK Ig2-like domain has two positively charged patches.

Immunoglobulins can contain glycans within the variable region [19]. Consensus amino acid motifs (N-X-S/T) for N-linked glycosylation sites can either be present within the germline sequence or can be acquired during somatic hypermutation. Enrichment of Fab glycans has been observed in autoimmunity [20, 21]. While conserved N-linked glycosylation sites are present in the antibody Fc regions, glycosylation of the variable domains is less common. Based upon their amino acid sequences, during affinity maturation all three mature MuSK mAbs **(Supplemental Figure 1)** were predicted to acquire or retain glycosylation sites within the variable regions [22], either in the heavy (MuSK1A and 3-28) or light chain (MuSK1B). The UCA of MuSK1B had an additional N-linked glycosylation site within the heavy chain that was lost during affinity maturation **(Supplemental Figure 1)**. Our electron density maps provided unambiguous experimental evidence that N81 of MuSK1A heavy chain and N20 of MuSK1B light chain were modified by N-linked glycosylation **(Figure 2 E and F)**. Carbohydrate residues could be modeled into additional electron density in MuSK1A at residue N81 of the heavy chain **(Figure 2 E)**, including the first two N-acetylglucosamine residues and a beta-D-mannose. In our mature MuSK1B structure, we were able to model the first N-acetylglucosamine residues **(Figure 2 F)**. Moreover, we recently confirmed these findings in an independent study (*Mandel-Brehm – BioRxiv 2020*).

We next sought to understand how affinity maturation could play a role in MuSK MG pathogenesis. Interestingly, the mature MuSK1A light chain, and to a lesser extent mature MuSK1A heavy chain, CDR loops were largely negatively charged **(Figure 2 G)**, while the germline sequence in the CDR regions may have more neutral or additional positive charge **(Figure 2 I)**. For example, G28D in L CDR1, Y101D in H CDR3, or K30N in L CDR1 might make this region more negatively charged, while K52M in L CDR2 might serve to neutralize positive charge present in the UCA sequence. Similarly, the mature MuSK1B CDR loops were largely negatively charged **(Figure 2 H)**, while sequence related Fabs to the UCAs tend to be largely positively charged **(Figure 2 I),** and hence the mutations from the UCA to the mature Fab might serve to increase the negative charge of the CDR regions. For example, the mutations S51D in L CDR2, and G32D and S57N respectively in H CDR1 and H CDR2, might all increase the negativity of the antigen binding site. Notably, the MuSK Ig1-like domain is predominantly negative, while the MuSK Ig2-like domain is highly positively charged [23] **(Figure 2 J)**. As the MuSK1A and MuSK1B epitopes have been mapped at the domain level to the MuSK Ig2-like domain [16], this might suggest that MuSK1A and MuSK1B likely bind to one or both of the basic patches on the Ig2-like domain of MuSK. In summary, these collective structural data indicate that the MuSK1A and MuSK1B Fabs share common structural features with Fabs of similar composition and include occupied variable region glycosylation sites. These findings further suggest that acquired mutations may strengthen the binding affinity for the basic patches on the MuSK Ig2-like domain, by altering the electrostatic interactions at the antigen-antibody interface.

### The pathogenicity of the MuSK autoantibodies correlates with both affinity and valency

We next sought to evaluate how the amino acid changes resulting from SHM contributed to pathogenicity. We previously demonstrated that all three mature mAbs are pathogenic when tested using an established *in vitro* AChR clustering assay [16]. Accordingly, we investigated whether the UCA counterparts of the mature mAbs disrupt AChR clustering in C2C12 myotubes. In addition to the divalent mAbs, we tested whether the amino acid substitutions from SHM had an impact on pathogenicity by testing the mature and UCA monovalent Fabs, which emulated Fab-arm exchange products.

C2C12 myotubes were incubated with Agrin, the neuronal ligand that stimulates AChR clustering, together with MuSK mAbs, MuSK Fabs and control antibodies. AChR clusters were visualized and the mean number of AChR clusters quantified. The number of AChR clusters that formed in response to Agrin alone was assigned a value of 100%, and the number of AChR clusters that formed in the presence of the antibodies were expressed relative to this value. The UCA of MuSK1A reduced the number of AChR clusters by 67.1% (p < 0.0001) and the UCA of MuSK1B by 73.0% (p < 0.0001) (**Figure 3 A and B**). Although the mature MuSK3-28 reduced the number of AChR clusters by 49.7% (p < 0.001), the UCA of MuSK3-28, unlike the other UCAs, failed to diminish the number of AChR clusters that formed in response to Agrin (% of agrin effect 98.3) (**Figure 3 B**). The Fabs from the mature MuSK1A (% of agrin effect 1.4) and MuSK1B antibodies (% of agrin effect 1.5) reduced AChR clustering to near background values (p<0.0001) (**Figure 3 B**). The UCA Fabs of MuSK1A and MuSK1B, however, had modest pathogenic capacity (AChR clustering reduced by ∼20% by each of them) (**Figure 3 B**). In contrast, the Fabs from the mature and UCA MuSK3-28 failed to inhibit Agrin-induced AChR clustering (**Figure 3 B**). In summary, the UCA of MuSK1A and MuSK1B mAbs retained their pathogenic capacity, but the UCA of the lower affinity mAb MuSK3-28 did not. However, in the monovalent configuration, which emulates Fab-arm exchange, only the mature, high affinity MuSK1A and MuSK1B antibodies were able to disrupt Agrin-induced AChR clustering.

**Figure 3.**
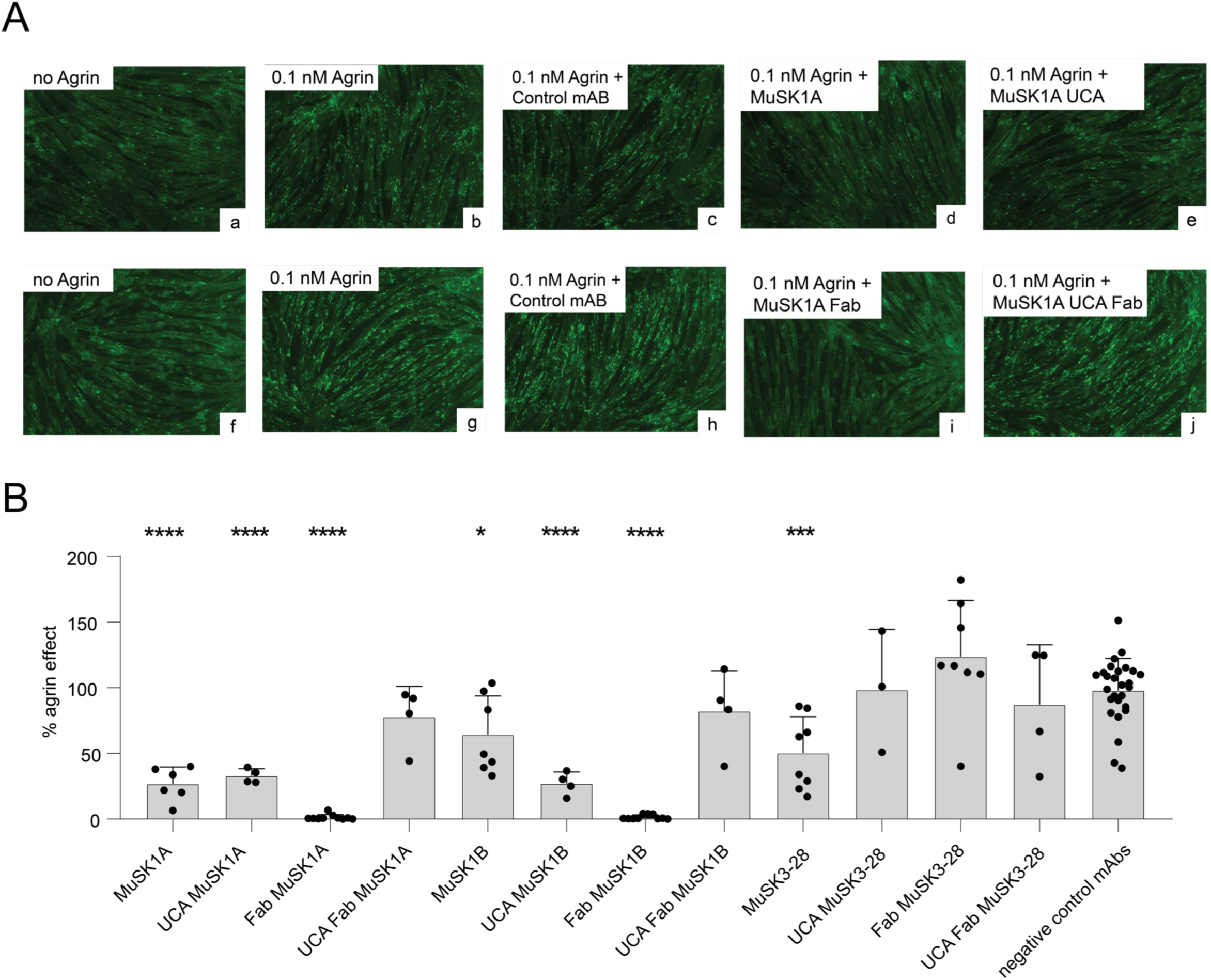
The unmutated common ancestors of MuSK autoantibodies have pathogenic capacity. AChR-clustering assay in C2C12 mouse myotubes demonstrates pathogenic capacity of MuSK mAbs. The presence of agrin in C2C12 myotube cultures leads to dense clustering of AChRs that can be readily visualized with fluorescent α-bungarotoxin and quantified. Pathogenic MuSK autoantibodies disrupt this clustering. The three different human MuSK-specific mAbs and their UCAs were tested for their ability to disrupt the AChR clustering. **(A)** Representative images (original magnification, ×100) from the clustering experiments are shown. (image a and f) Cultured myotubes do not show AChR clustering until (image b and g) agrin is added (bright spots reveal AChR clusters). The mAbs MuSK1A (d), MuSK1A UCA (e), and MuSK1A Fab (i) added at 1 μg/mL inhibit clustering. A Control mAb (c and h) and the MuSK1A UCA Fab (j) do not inhibit the formation of AChR clusters. **(B)** The effect of the mAb on Agrin dependent clustering was tested. Quantitative measurements of the C2C12 clustering were normalized to the Agrin-only effect of each individual experiment. Each data point represents the mean value from 2-8 individual values from in total 4-10 independent experiments. Bars represent the mean of means and error bars SDs. Multiple comparisons ANOVA (against the pooled results for the three human non-MuSK-specific mAbs), Dunnett’s test; * p<0.05, ** p<0.01, *** p<0.001, **** p<0.0001, only shown when significant.

## Discussion

Autoantibody-producing B cells are rare within the large circulating B cell repertoire. Given their direct and indispensable role in the pathology of many autoimmune diseases, the study of these B cells can provide unprecedented mechanistic insight. However, identifying and isolating these rare individual self-reactive B cells remains a challenge; accordingly, a limited number of studies have been reported. Such challenges notwithstanding, to further understand the mechanisms underlying autoimmune pathology, we recently identified and isolated a set of B cells from MuSK MG patients. We expressed whole recombinant human monoclonal antibodies from these cells, then confirmed their specificity and pathogenic capacity [15, 16]. In the current study, we used these mAbs to investigate both the origin and evolution of the autoantibody response in MuSK MG.

We reverted the accumulated somatic mutations of the mAbs back to their germline configuration, thus generating UCAs. The UCAs served as a proxy for the divalent B cell receptor expressed by naïve B cells. This afforded us the opportunity to test whether the naïve B cells were initially stimulated by the self-antigen, MuSK. We also expressed whole IgG and their Fabs so that we could test the contribution of somatic hypermutation to both monovalent- and divalent-binding, thus emulating the products of Fab-arm exchange from IgG4 antibodies. With this array of autoantibody constructs, we tested binding to the autoantigen, pathogenic capacity and the affinity between the autoantibody and the cognate self-antigen. We found that the UCAs of MuSK autoantibodies retain remarkable binding to MuSK. The endogenous heavy and light chain combination was required for this binding. The binding was not a product of polyspecificity, and the UCAs recognized the same domain on MuSK as the mature mAbs, suggesting that naïve B cells are initially activated by MuSK and that MuSK drives SHM. The UCAs retained the pathogenic capacity of the parental mature mAbs, but only when they were presented to the antigen as dimeric mAbs and not as monovalent Fabs. Affinity measurements demonstrated that SHM was a critical requirement for pathogenic capacity of the monovalent Fabs. Structural data indicated that, as SHM led to a large increase in binding affinity, it also resulted in an increasingly negatively charged antigen binding surface on both MuSK1A and MuSK1B. Due to the largely positively charged surface on MuSK Ig2-like domain, it is tempting to speculate that this charge complementarity contributed to the increased binding affinity. Interestingly, a reversion mutation for A25D near H CDR1 in the FR1 of MuSK3-28 largely abolished MuSK3-28 binding to MuSK in our CBA **(Supplemental Figure 3 A)**. Homology modeling of MuSK3-28 (data not shown) suggests that introducing this negatively charged residue during affinity maturation, which is buried in the FR region, could alter the conformation of the H CDR1 loop and may partially explain the observed difference in the k_on_ observed between the mature and UCA MuSK3-28 Fab. These collective data would suggest that, pathology occurs when a high affinity threshold, governed by a slow off-rate, is reached by a Fab-arm exchanged IgG4.

Our findings provide insight regarding two fundamental aspects of MuSK MG immunopathology. First, the MuSK autoantigen appears to initiate the response with germline encoded mAbs serving as a proxy for the BCR on the surface of naïve B cells. In a number of autoantibody-driven autoimmune diseases, there is speculation, as well as some empirical support, for exogenous antigen as the initiator and propagator of the autoimmune response. This does not appear to be the case for MuSK MG. Secondly, our data show that SHM is a requirement for monovalent IgG autoantibodies (the product of Fab-arm exchange) to bind MuSK, and to reach pathogenic capacity.

### Autoreactivity in the naïve repertoire

Our findings indicate that the B cell receptors (BCR) expressed on naïve B cell clones can bind to MuSK. B cells expressing autoreactive BCRs are normally eliminated from the maturing repertoire by mechanisms present at two distinct checkpoints along the B cell development pathway, thereby reducing the development of immune responses against self-antigens [15, 18]. A number of autoimmune diseases [24], including both AChR and MuSK MG [25], include defective B cell tolerance checkpoints. Consequently, the repertoire includes a higher proportion of self-reactive B cells than found in healthy (non-autoimmune) individuals.

Given that the self-antigen, MuSK, bound to germline encoded mAbs - emulating the BCR of naïve B cell clone - it is reasonable to speculate that MuSK is both initiating and driving the autoimmune response. This finding is unusual, as similar investigations of germline encoded human mAbs demonstrate a remarkable lack of autoantigen binding that is, conversely, robust in their mature counterparts. Unmutated revertants of monoclonal autoantibodies from patients with pemphigus vulgaris and systemic lupus erythematosus do not bind self-antigen [13, 26, 27]. Similar patterns were found with unmutated revertants of anti-cytokine autoantibodies in AIRE-deficient patients [28] and pulmonary alveolar proteinosis patients [29]. These findings contrast with those acquired with antibodies that develop toward exogenous (non-self) antigens during a normal immune response. Examples includes viral infections [30–33] and responses to influenza hemagglutinin, wherein unmutated revertants of virus antigen-specific monoclonal antibodies retain binding activity. These scenarios, however, must be carefully considered, as there are some exceptions of autoantibody germline binding [34, 35]. The collective findings may point toward limitation of the approaches available to measure antibody-antigen interactions, including ELISAs and live cell-based assays, neither of which may be sufficiently sensitive to measure low affinity interactions or accurately emulate the antigen/autoantigen binding to the BCR of naïve B cell in situ, such as lymphoid tissue.

MuSK appears to both initiate and propagate the B cell response, which is a rare finding in human autoimmunity [35]. The generation of pathogenic autoantibodies may be the result of an aberrant immune response to self-antigen that is both initiated and driven by the same self-antigen. While alternative models of autoimmunity suggest that exogenous antigen can be the initiators of an immune response that then evolves to autoimmune through other mechanisms such as molecular mimicry. While corroborating empirical validation remain to be acquired for such cross-reactive mechanisms, there are several intriguing examples. This list includes a variant of pemphigus in which monoclonal autoantibodies to the target autoantigen (desmoglein 1) also bind to LJM11, a salivary protein from a sand fly known to transmit a co-morbid disease [36]. And also, in autoimmune NMO there is evidence that T cells, which drive B cells to produce pathogenic autoantibodies, are initiated by a high frequency of specific gut bacteria (*Clostridium perfringens*) [37, 38]. Although, we cannot exclude whether another antigen with high epitope similarity to the MuSK Ig2-like domain exists and is the basis for the affinity maturation in MuSK MG, we show here that MuSK itself appears to both initiate and propagate the process. These findings are consistent with the presence of defective counterselection, as this fundamental aspect of autoimmunity may contribute to the reservoir of self-reactive naïve B cells, including those that served as precursors to the pathogenic autoantibodies we isolated from the MuSK MG patients.

### Relevance to disease mechanisms

The MuSK specific mAbs that we generated were derived from two IgG4-expressing B cells and one IgG3 expressing B cell [15, 16]. MuSK specific mAbs have also been derived from IgG1-expressing B cells [14]. Neither IgG1 nor IgG3 participate in Fab-arm exchange. However, the collective findings regarding the pathogenic capacity of MuSK autoantibodies demonstrate that they require both high affinity, as described here, and monovalent binding [14] for robust pathogenic capacity. Thus, the roles of IgG1 and IgG3 MuSK autoantibodies are not clear. Low titers of autoantibodies with IgG1 and IgG3 subclass were described in a cohort of patients with high titers for IgG4 antibodies [39–41], so the pathology of MuSK MG may be a response to both IgG1-3 and IgG4 autoantibodies. Both IgG1 and IgG3 subclasses are effective at initiating the complement cascade, while IgG4 does not [42]. It is not known whether these IgG1 and IgG3 subclasses bind to MuSK at the neuromuscular synapse in humans, and if so, whether complement-mediated damage contributes to disease.

There is no reported deposition of complement at the NMJ in MuSK MG patients as it has been in AChR MG [43]. MuSK mAbs do demonstrate pathogenic capacity using *in vitro* AChR clustering assays, but they are not as effective as their monovalent counterparts. In addition, the divalent antibodies stimulate rather than inhibit MuSK phosphorylation, whereas their monovalent counterparts, like IgG4 antibodies in sera from MuSK MG patients, inhibit MuSK phosphorylation [14, 16, 44]. The difference between the divalent and monovalent antibodies is likely due to the dual activity of the divalent antibodies, as the divalent antibodies can dimerize MuSK, known to stimulate transphosphorylation [45], and at the same time inhibit binding of Lrp4 to MuSK. Experiments that test the pathogenic capacity of the divalent and monovalent mAbs *in vitro* and in rodent models will likely be required to gain further understanding into the role of the different subclasses in disease. Finally, the ability to stimulate MuSK phosphorylation suggest that mAbs with these properties may have therapeutic value, especially if their ability to stimulate complement is attenuated.

The remarkably high affinity of these anti-MuSK autoantibodies offers additional insight into the mechanisms of immunopathology. The affinity has only been described for a small number of pathogenic human autoantibodies; in contrast to many reported avidity measurements or estimates of avidity, which are derived from divalent mAbs. The association and dissociation rates need to be separately measured to calculate the true affinity or avidity. Measuring binding over a broad range of concentrations can give an estimate of the binding to an antigen, and as such is often used as an affinity/avidity estimate. An AChR mAb bound with an estimated avidity in the sub-nanomolar range [46], and mAbs binding AQP4, derived from patients with NMO, have estimated avidities covering a broad range, from 24 nM to 559 nM [24]. The affinity measurements of a human mAb against respiratory syncytial virus (Fab19) and several reversion intermediates, together with the germline-encoded Fab, showed that SHM increased affinity and resulted in a faster association rate that was correlated to an increase in antiviral neutralizing activity [47], which was similar to earlier findings on the binding kinetics of an immune response [48]. Two of the mAbs described here have affinities for MuSK in the sub-nanomolar range, which was driven by a remarkably slow off-rate. Germline versions displayed diminished affinity with a much faster off-rate, which was correlated with diminished pathogenicity. Thus, these autoantibodies, once they have engaged MuSK at the neuromuscular synapse, may remain bound to MuSK for an extended period of time, which may be a key property of disease. High-affinity, monovalent binding also demonstrates that cooperative binding of both antibody arms (avidity) is not required. These findings shed light on the non-essential role of the non-MuSK binding arm of a Fab-arm exchanged autoantibody. Nonetheless, it is reasonable to speculate that the dual specificity of a Fab-arm exchanged antibody may be a property of its pathogenic capacity. Engineered bispecific anti–HIV-1 antibodies (BiAbs) that bind to two separate epitopes on the virus are superior in viral neutralization compared with their parental antibodies [49]. In a similar scenario Fab-arm exchanged IgG4 with a single anti-MuSK arm may increase or adopt effective binding capabilities through interaction of the opposite arm with a second component of the neuromuscular junction. Our data suggest that this model may not be sustainable given that affinity of the single-arm MuSK mAb is remarkably high and pathogenic.

### Immunomodulating therapy in MuSK MG

The mechanistic details presented here also provide support for both current biologic treatment strategies and new therapeutic targets to consider. First, they offer additional justification for the use of B cell depletion. Rituximab (RTX), which affects B cell depletion through targeting CD20, is highly beneficial in treating MuSK MG [50, 51]. CD19-specific inebilizumab targets a broader range of B cell subsets. The use of inebilizumab has been demonstrated to successfully treat NMO [52]. Precise targeting of MuSK autoantibody-expressing B cells with CAAR-T cells [53] gains further justification by the additional mechanistic details provided in the current study. An agonist antibody to MuSK, which stimulated muscular signaling, showed promising effects in a mouse model of amyotrophic lateral sclerosis (ALS) [54]. Our data, showing that early naïve precursors of antibody-secreting cells (ASCs) have high affinity BCRs, provides further interest in the use of such technology, as naïve precursors, as well as ASCs may be eliminated by this strategy. Our findings also support the perception that complement inhibitors are unlikely to have a beneficial effect in MuSK MG, unless the IgG1-3 antibodies participate in disease and engage complement at the synapse. Newer therapeutic approaches for treating autoantibody-mediated disease seek to target circulating autoantibodies by inhibiting the neonatal FcR, resulting in diminished autoantibody titers [55]. Given the requirement for high affinity Fab-arm exchanged autoantibodies in MuSK MG opens the possibility that inhibition of Fab-arm exchange may represent a viable strategy. However, the biological relevance of Fab-arm exchange in the normal immune response is not well-understood. Thus, ‘off-target’ consequences should be considered.

### Limitations

Our study is, of course, not without limitations. First, the process of producing UCA versions of mature antibodies is not absolute without having the germline B cell clone in-hand. We made every effort to determine whether the CDR3 region harbored somatic mutations, including reverting the D-gene segment and using a statistical model that evaluated proposed germline configurations [56]. Although not absolute, our reversions are as thorough as experimentally possible and thus provide the best approximate representation of the true germline BCR.

While human mAbs provide an exceptional opportunity to gain insight into the mechanism of B cell mediated autoimmunity, their biological relevance requires careful interpretation. Most important in this regard is how well a B cell–derived mAb represents both the circulating repertoire of autoantibodies and the repertoire at the site of the antigen and tissue injury. The mAbs studied here were derived from plasmablasts and memory B cells. While the former provides some assurance of expression, given their antibody-secreting phenotype, which actively contributes to circulating Ig, the same cannot be said of memory B cells. Coupling of BCR sequencing with a mass spectrometry-derived Ig proteome will be required to more accurately link and identify which autoreactive B cells contribute to circulating autoantibodies in MuSK MG [57].

### Speculative mechanism of MuSK MG immunopathology

The collective findings of this study provide new insights into the mechanism of pathogenic autoantibody production in MuSK MG. In a previous study, we demonstrated that MuSK MG, like many autoimmune diseases, includes defective B cell tolerance checkpoints [25]. The consequence of which is a naïve B cell repertoire populated with clones that circumvented counterselection. In the current study, we show that the MuSK MG patient naïve B cell repertoire includes clones that are capable of binding to MuSK with high affinity. We suggest that such clones would not have escaped counterselection in the presence of well-functioning B cell tolerance checkpoints. Thus, these autoreactive naïve B cells bind self-antigen and participate in initiating B cell differentiation toward memory B cells and ASCs that contribute to disease. Our study further suggests that this process is driven throughout by MuSK. At the stage of autoantibody production, somatic hypermutation is required to generate autoantibodies of extraordinarily high avidity. Fab-arm exchange then takes place and functionally monovalent autoantibodies that have exceeded a high affinity threshold bind to MuSK at the neuromuscular synapse and cause disease. Targeting of B cells, B cell maturation or Fab-arm exchange represent viable therapeutic strategies for MuSK MG treatment based on this model.

## Materials and Methods

### Autoantibody variable region site directed mutagenesis

To identify the base changes that arose through affinity maturation process, the sequences of the MuSK mAbs were aligned against the 2018-02-24 germline reference set with IMGT/HighV-QUEST v1.6.0 (the international ImMunoGeneTics information system^®^) [58]. The mAbs were reverted to the UCA sequence in a step-by-step manner using the Q5^®^ Site-Directed Mutagenesis Kit (NEB) according to manufacturer’s instructions. The primers were designed with NEBaseChanger.

### Recombinant expression of MuSK, human monoclonal antibodies (mAbs) and Fabs

For crystallography, MuSK1A was expressed in Sf9 cells using the baculovirus expression system. The heavy and light chain variable domains of MuSK1A were fused to a standard human IgG1 heavy and light chain constant domain and cloned into pFastBacDual to create pBE1719. A C-terminal 6xHis-Tag was included in the heavy chain for subsequent use in purification. Sf9 cells were grown in suspension in Sf-900 II media for three days before harvesting. Clarified Sf9 supernatants were filtered and MuSK1A was purified by immobilized metal affinity chromatography (IMAC) (Ni Sepharose excel, GE Life Sciences) followed by size exclusion chromatography (Superdex 200 Increase 10/300 GL column) on an ÄKTA pure system in 20 mM Tris pH 8.0, 150 mM NaCl. Both the variable and constant domain of the light chain of MuSK1B was cloned into pCDNA3.4-TOPO vector to create pBE1779 for protein expression for crystallography of MuSK1B. The variable domain of the heavy chain of MuSK1B was fused to a standard human IgG1 CH1 constant domain and subsequently cloned into the pCDNA3.4-TOPO vector to create pBE1775. MuSK1B was then expressed from Expi293F cells transfected using ExpiFectamine with a 1:2 ratio of the corresponding heavy (pBE1775) and light chain plasmid (pBE1779). Culture supernatants were harvested seven days post-transfection and clarified by centrifugation. MuSK1B was purified from culture supernatants using HiTrap Protein G High Performance column (GE Life Sciences) equilibrated in 20 mM Tris pH 8.0, 150 mM NaCl. Protein was eluted in 0.1 M glycine-HCL, pH 2.5 and the pH was immediately neutralized with 1 M Tris pH 9.0. MuSK1B was then subsequently purified using size-exclusion chromatography (Superdex 200 Increase 10/300 GL column) on an ÄKTA pure system in 20 mM Tris pH 8.0, 150 mM NaCl.

For affinity measurements, the ectodomain construct of MuSK (ectoMuSK) (aa1-472) was expressed in Sf9 cells using a baculovirus system and was designed with a C-terminal Avitag and 6xHis-tag. Clarified Sf9 culture supernatants were filtered and ectoMuSK was purified by IMAC (HisTrap Excel, GE Life Sciences), and exchanged into low-salt TBS for site-specific biotinylation. ectoMuSK was incubated with BirA in the presence of ATP, magnesium acetate, and D-biotin for 4 hours at room temperature, then purified by size-exclusion chromatography (Superdex 200 Increase 10/300 GL).

All UCA Fab expression constructs that were used for affinity measurements and the AChR clustering assay were cloned in a pcDNA3.4-TOPO vector for expression and secretion from mammalian cells. Expi293F cells were transiently transfected using ExpiFectamine and culture supernatants were harvested 5 days post-transfection and clarified by centrifugation. Clarified Expi293F culture supernatants were filtered and Fabs were purified by successive affinity and size-exclusion chromatography (on HiTrap Protein G and Superdex 200 Increase 10/300 GL columns, respectively, from GE Life Sciences) using an AKTA pure system.

The mAbs were produced as previously described [16]. Briefly, HEK293A were transfected with equal amounts of the heavy and the corresponding light chain plasmid using linear PEI (Polysciences Cat# 23966). The media was changed after 24 h to BASAL media (50% DMEM 12430, 50% RPMI 1640, 1% antibiotic/antimycotic, 1% Na-pyruvate, 1% Nutridoma). After 6 days the supernatant was harvested and Protein G Sepharose® 4 Fast Flow beads (GE Healthcare) were used for antibody purification. For expression of mAbs using the human IgG3 constant region, a vector containing human IgG3 was purchased from Addgene (pVITRO1-102.1F10-IgG3/λ) and then cloned into a vector for recombinant IgG expression that we previously engineered [59].

### Affinity measurements

Fab binding to MuSK was measured by biolayer interferometry using an Octet RED96 system (ForteBio). Biotinylated ectoMuSK was immobilized on streptavidin coated sensors and incubated with varying concentrations of purified Fabs. During the experiment, the reaction plate was maintained at 30 °C and shaken at 1000 rpm. Data were fit using the Octet System data analysis software. After subtraction of signal from a reference sensor loaded with ectoMuSK and incubated with buffer, response curves were aligned at the equilibration step. Inter-step correction was applied to align association and dissociation curves and Savitzky-Golay filter was applied to remove high-frequency noise. Processed association and dissociation curves were fit globally using a 1:1 binding model to obtain kinetic constants for each Fab.

### Live cell-based autoantibody assay

HEK293T (ATCC^®^ CRL3216™) cells were transfected with either full-length MuSK-GFP (provided by David Beeson and Patrick Waters, University of Oxford) or different ectodomain variants of MuSK-GFP (previously described in [16]). On the day of the CBA, the antibodies were added to the transfected cells in either a dilutions series (10 – 0.02 µg/ml) or at a concentration of 10, 1 or 0.1 µg/ml. For the epitope determination assay all constructs were measured at 10 µg/ml. The binding of each mAb was detected with Alexa Fluor^®^-conjugated AffiniPure Rabbit Anti-Human IgG, Fcγ (309-605-008, Jackson Immunoresearch) on a BD LSRFortessa^®^ (BD Biosciences). FlowJo software (FlowJo, LLC) was used for analysis.

### Crystallization and structure determination

Gel filtration fractions containing purified MuSK1A were concentrated to 10 mg/ml. MuSK1A crystals were grown at 18 °C by vapor diffusion and after ≈ 4 days grew from drops consisting of 100 nL protein plus 100 nL of a reservoir solution consisting of 20% w/v PEG 6K (Precipitant) and 0.1 M BICINE 8.5 pH (Buffer) from the JCSG Core I (Qiagen) screen. Reservoir solution was supplemented with 15% ethylene glycol for cryoprotection. Gel filtration fractions containing purified MuSK1B were concentrated to 10 mg/ml. MuSK1B crystals grew after 1 day at 18 °C by vapor diffusion from drops consisting of 100 nL protein plus 100 nL of a reservoir solution consisting of of 0.085 M HEPES pH 7.5, 17% PEG 4K, 8.5% Isopropyl alcohol, and 15% Glycerol from the JCSG Core II (Qiagen) screen. Reservoir solution was supplemented with 5% glycerol for cryoprotection. For both MuSK1A and MuSK1B, native diffraction data was collected at GM/CA-CAT beamline 23-ID-B at the Advanced Photon Source and reduced using XDS **(Supplemental Table 2)** [60]. MuSK1A was indexed to P2_1_ and MuSK1B was indexed to P2_1_2_1_2_1_. Crystallographic data quality was assessed for outliers and the potential presence of twinning using Xtriage [61], and the high-resolution limit of each dataset was assessed using CC1/2 and cut where this metric dropped below ∼50% in the highest resolution shell, corresponding to CC* values of 0.725 and 0.781 for the MuSK1A and MuSK1B models, respectively, after refinement [62]. The MuSK1A asymmetric unit (ASU) consists of two sets of MuSK1A Fabs, and MuSK1A was phased by molecular replacement using Phaser [63] first using the Fab constant domains from PDB code 6DW2 as a search model, and then using the Fab variable domains from 6DW2 as a search model. MuSK1B has one Fab in the ASU and molecular replacement was carried out using the autoMR module from Phaser. First the Fab constant domains from PDB code 7FAB were used as a search model, followed by the Fab variable domains from PDB code 8FAB. The resulting models were adjusted in Coot [64] and refined using Phenix [65–67]. Protein models were validated using MolProbity [68].

### AChR clustering assay

The C2C12 AChR clustering assay was performed essentially as reported [16]. AChR clusters were counted using ImageJ software. For each condition duplicate wells were used and the mean of the clusters per visual field per condition were calculated. Experiments were performed at a minimum of three repetitions and were normalized for the effect of Agrin. Reported results are from experiments in which a minimum three-fold effect of Agrin-induced clustering over the baseline was observed.

### Statistics

Statistical significance was assessed with Prism Software (GraphPad; version 8.0) by two-tailed unpaired t-test or multiple comparison ANOVA with Dunnett’s correction for all flow cytometry-based assays and multiple comparison ANOVA with Dunnett’s correction for AChR clustering on the C2C12 cells.

## Acknowledgements

The authors thank Karen Boss for providing editorial assistance.

## Author contributions

This study was originally conceived, then initiated and directed by KCO. MLF led the laboratory work at Yale; designing, executing and interpreting experiments associated with the mutagenesis, mAb expression, C2C12 assay, and the CBA. MLF directed PAS in performing and validating the mutagenesis and expressing the mAb variants. DCE and CV planned, designed and executed the crystal structures. CV, LK, and RR produced the Fabs and executed the affinity measurements. KT helped designing and executing the mutagenesis and CBAs. PS assisted with C2C12 measurements. RJ executed the bioinformatic analyses. DCE and SJB provided key contributions to the overall project scope, experimental design and directed the laboratory work at NYU. RJN provided the clinical specimens from which the mAb were derived and provided insight on the clinical and therapeutic relevance of the findings. The manuscript was initially drafted by MLF and KCO. All authors contributed to the editing and revising.

## Graphical abstract

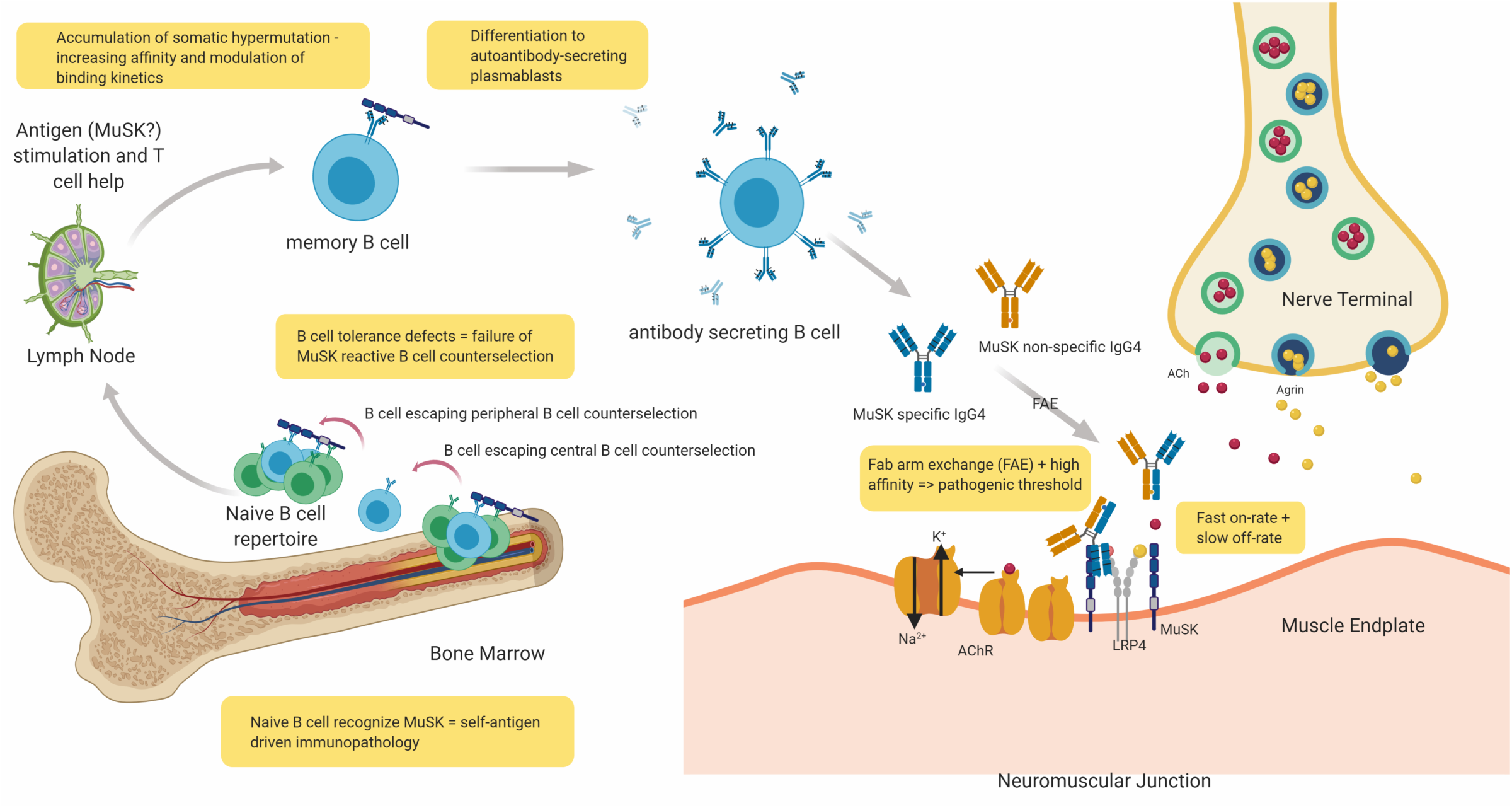

## Supplemental materials

**Supplemental Table 1.**
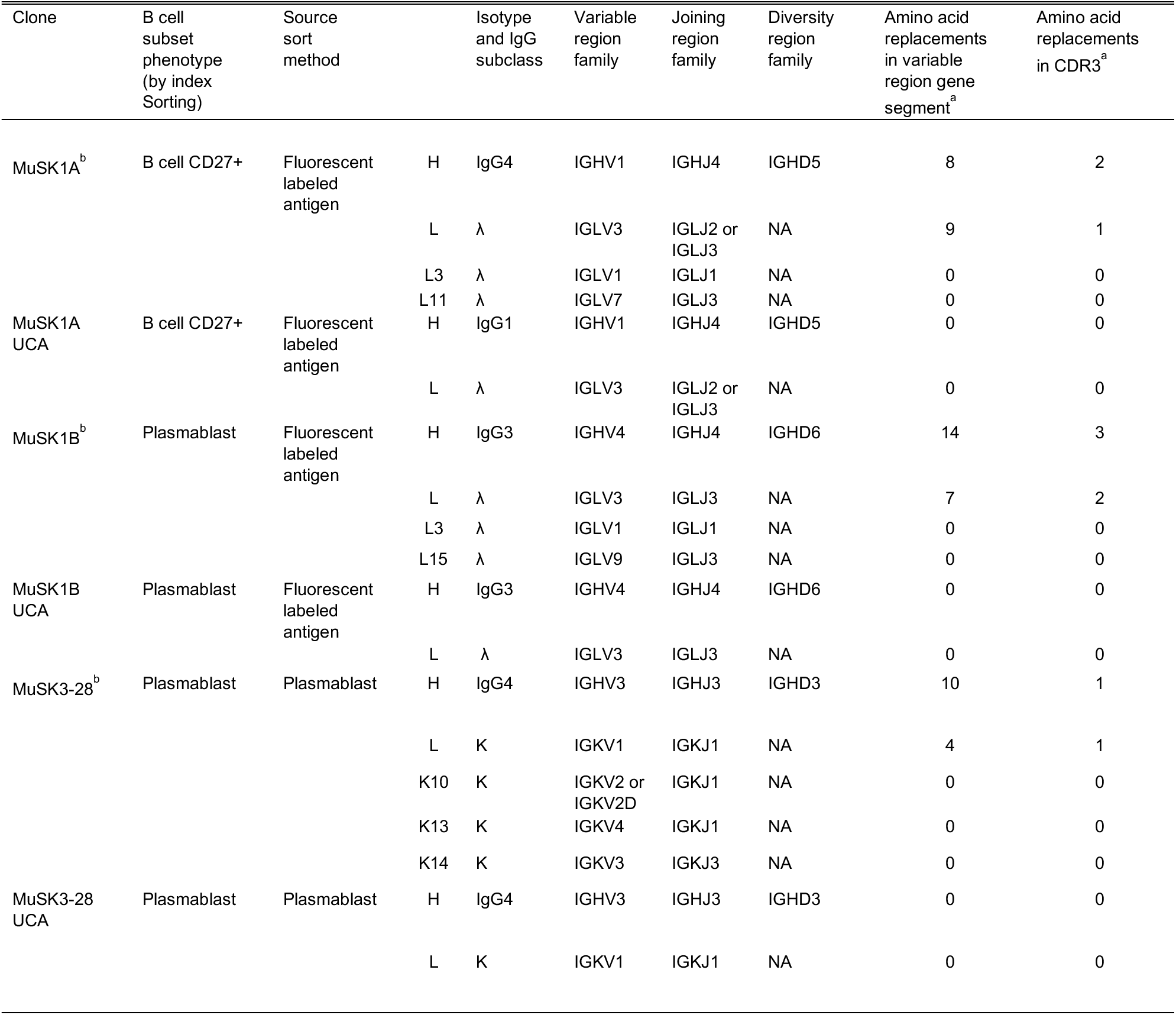
Molecular characteristics of MuSK-binding human recombinant mAbs. The molecular characteristics for the mature and UCA sequences of each antibody are shown. Additionally, the different light chains that were used for the light chain switch experiments are given **(Supplemental Figure 6)**. The replacement mutations in the variable region gene segment were counted from the beginning of framework 1 through the invariable cysteine at position 104. The mutations in the CDR3 were counted between cysteine 104 and the invariable tryptophan (W) or phenylalanine (F) at position 118 in both the heavy chain and the light chain respectively. No FR4 mutations were observed. These mAbs had been produced and studied in our previous investigations: MuSK1A and MuSK1B (Takata, Stathopoulos et al. 2019) and MuSK3-28 (Stathopoulos, Kumar et al. 2017).

**Supplemental Table 2.**
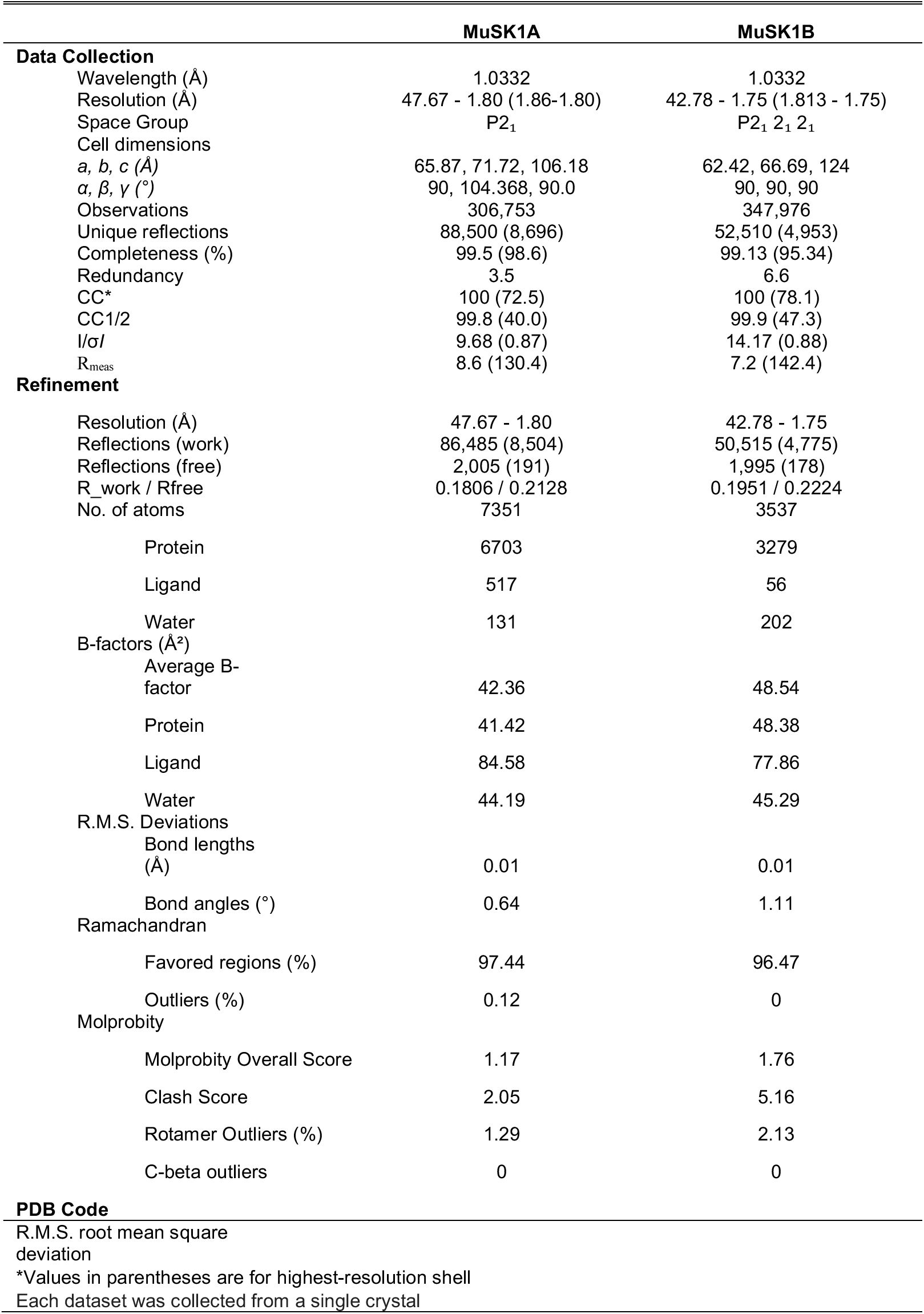
Crystallographic data collection and refinement statistics, related to MuSK1A and MuSK1B, related to Figure 2.

**Supplemental Figure 1.**
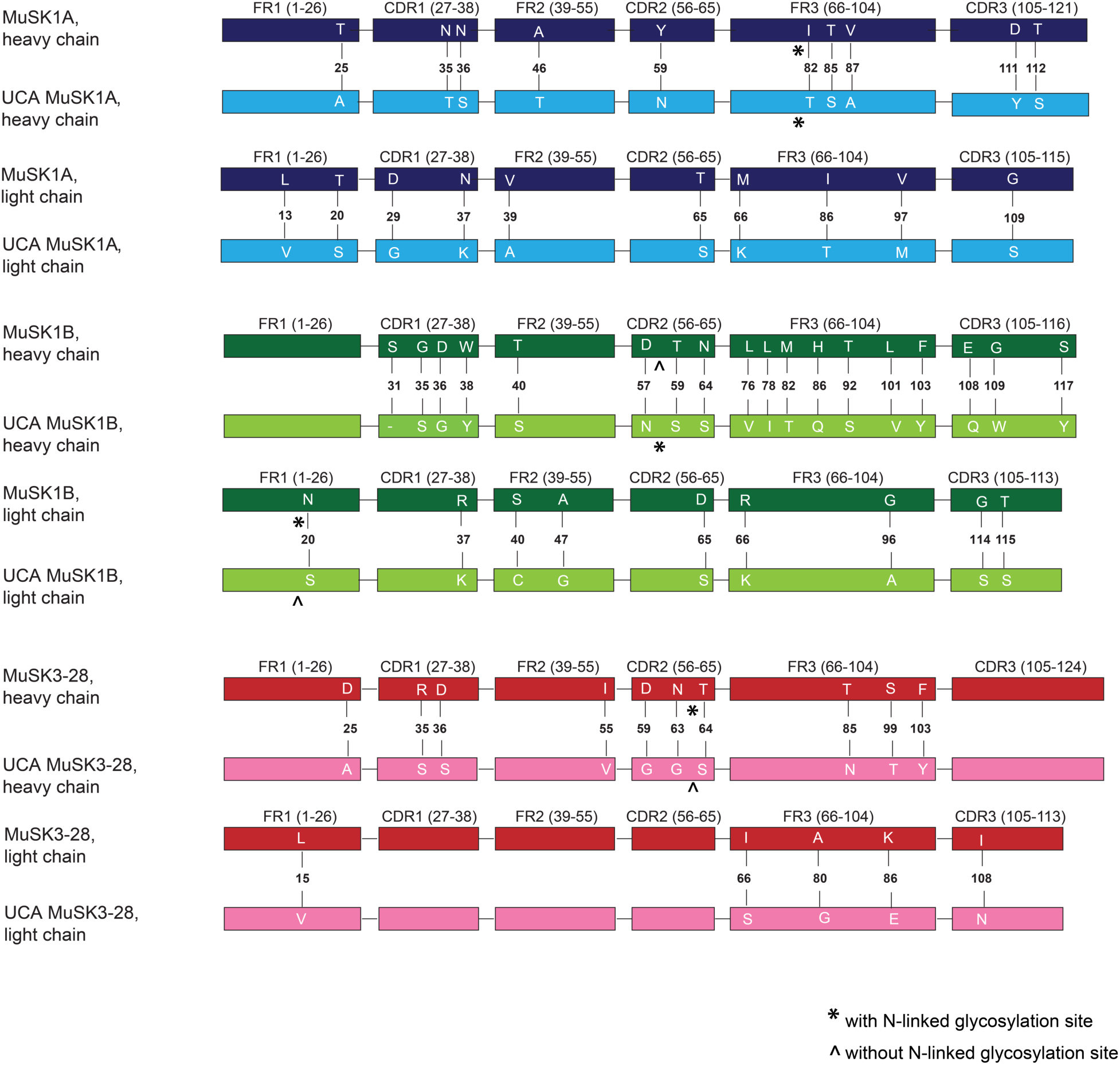
Comparison of the heavy and light chain of the MuSK mAb to their UCA sequence. Illustration of the differences in amino acid (AA) sequence between the mature mutated heavy and light chains to the UCA sequences. The different AA are shown in white letters. The number indicates the position of the AA within the variable region. The glycosylation sites were indicated with * if present and ^ if not present.

**Supplemental Figure 2.**
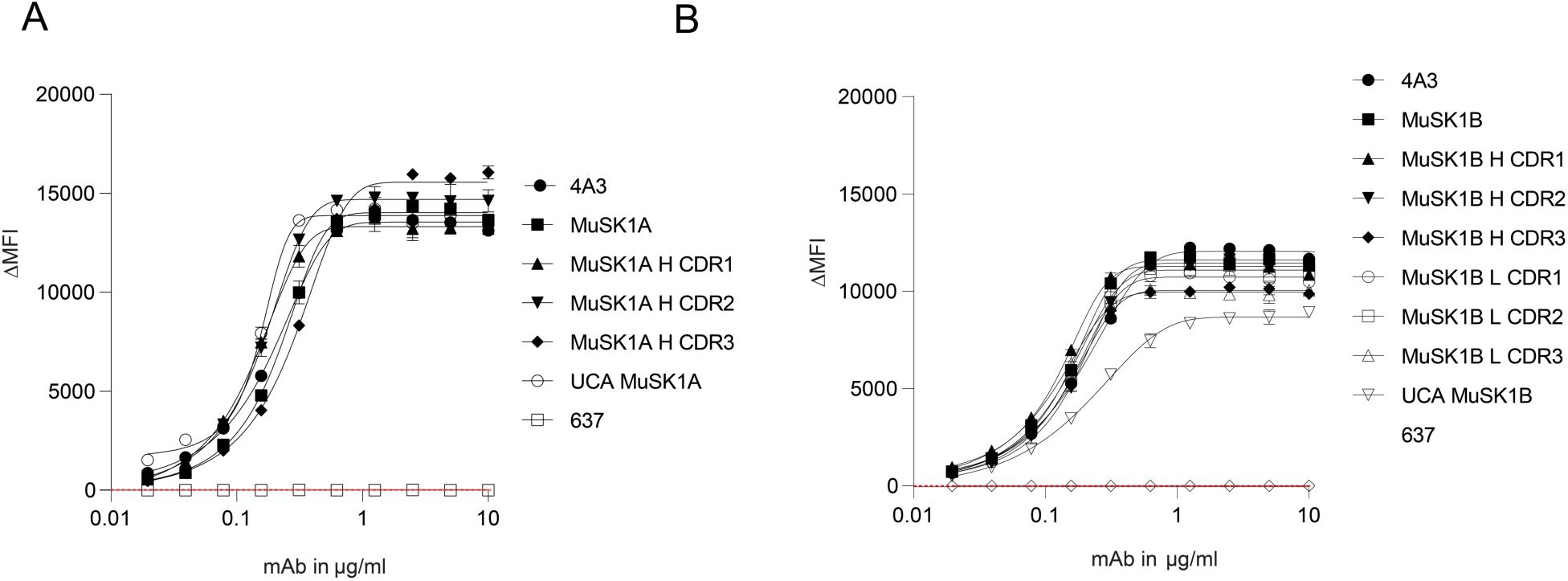
The intermediates of MuSK1A and MuSK1B bind to the MuSK autoantigen. MuSK-specific mAbs were tested for surface binding to MuSK on MuSK-GFP–transfected HEK cells. **(A and B)** Binding to MuSK was tested over ten two-fold serial dilutions of MuSK1A **(A)** and MuSK1B **(B)** ranging from 10 – 0.02 µg/ml. Humanized MuSK mAb 4A3 was used as the positive control and AChR-specific mAb 637 as the negative control. The ΔMFI was calculated by subtracting the signal from non-transfected cells from the signal of transfected cells. Each data point represents a separate replicate within the same experiment. Symbols represent means and error bars SDs. Values greater than the mean + 4SD of the negative mAb at 1.25 µg/ml (indicated by the horizontal dotted line) were considered positive.

**Supplemental Figure 3.**
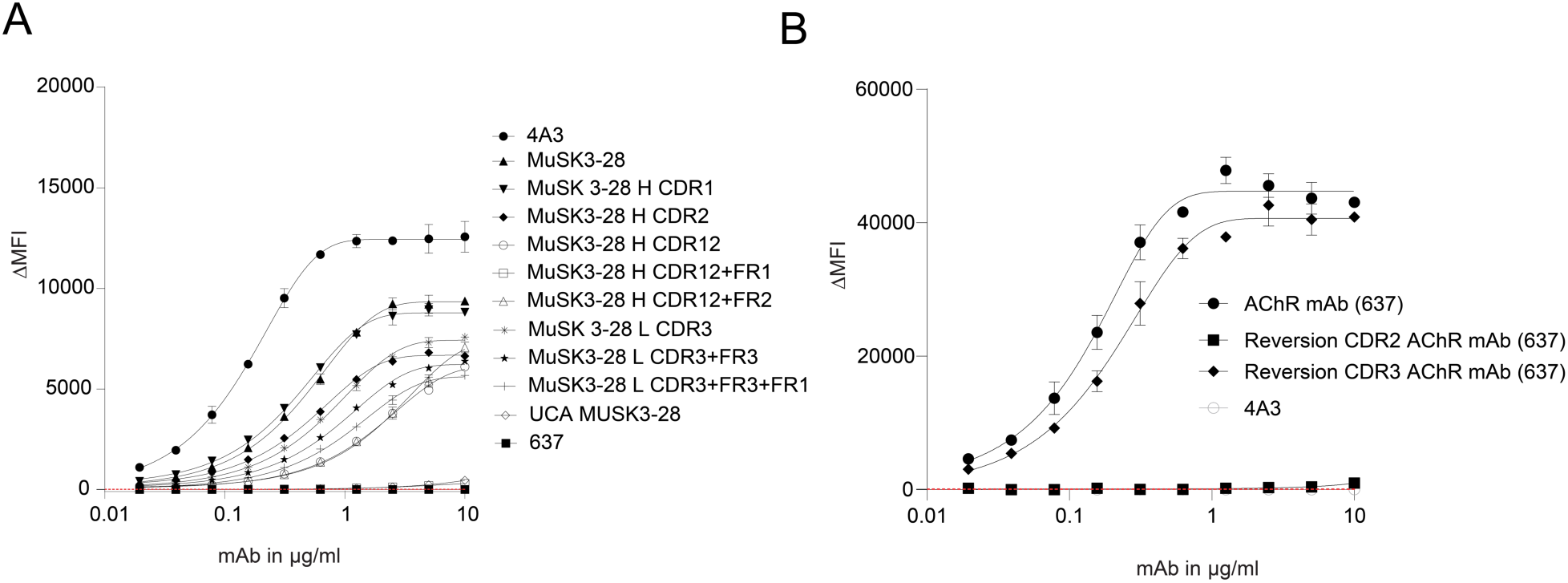
Somatic hypermutation contributes to the antigen binding of MuSK mAbs and AChR mAb 637. MuSK-specific mAbs were tested for surface binding to MuSK on MuSK-GFP– transfected HEK cells and AChR-specific mAbs were tested for surface binding to AChR on AChR Rapsyn-GFP–transfected HEK cells. **(A and B)** Binding to MuSK **(A)** and AChR **(B)** was tested over ten two-fold serial dilutions of each antibody ranging from 10-0.02 µg/ml. Humanized MuSK mAb 4A3 was used as the positive control for MuSK and negative control for AChR and AChR-specific mAb 637 as the negative control for MuSK and positive control for AChR. The ΔMFI was calculated by subtracting the signal from non-transfected cells from the signal of transfected cells. Each data point represents a separate replicate within the same experiment, which was measured in duplicate. Bars represent means and error bars SDs. Values greater than the mean + 4SD of the negative mAb (637) at 1.25 µg/ml (indicated by the horizontal dotted line) were considered positive.

**Supplemental Figure 4.**
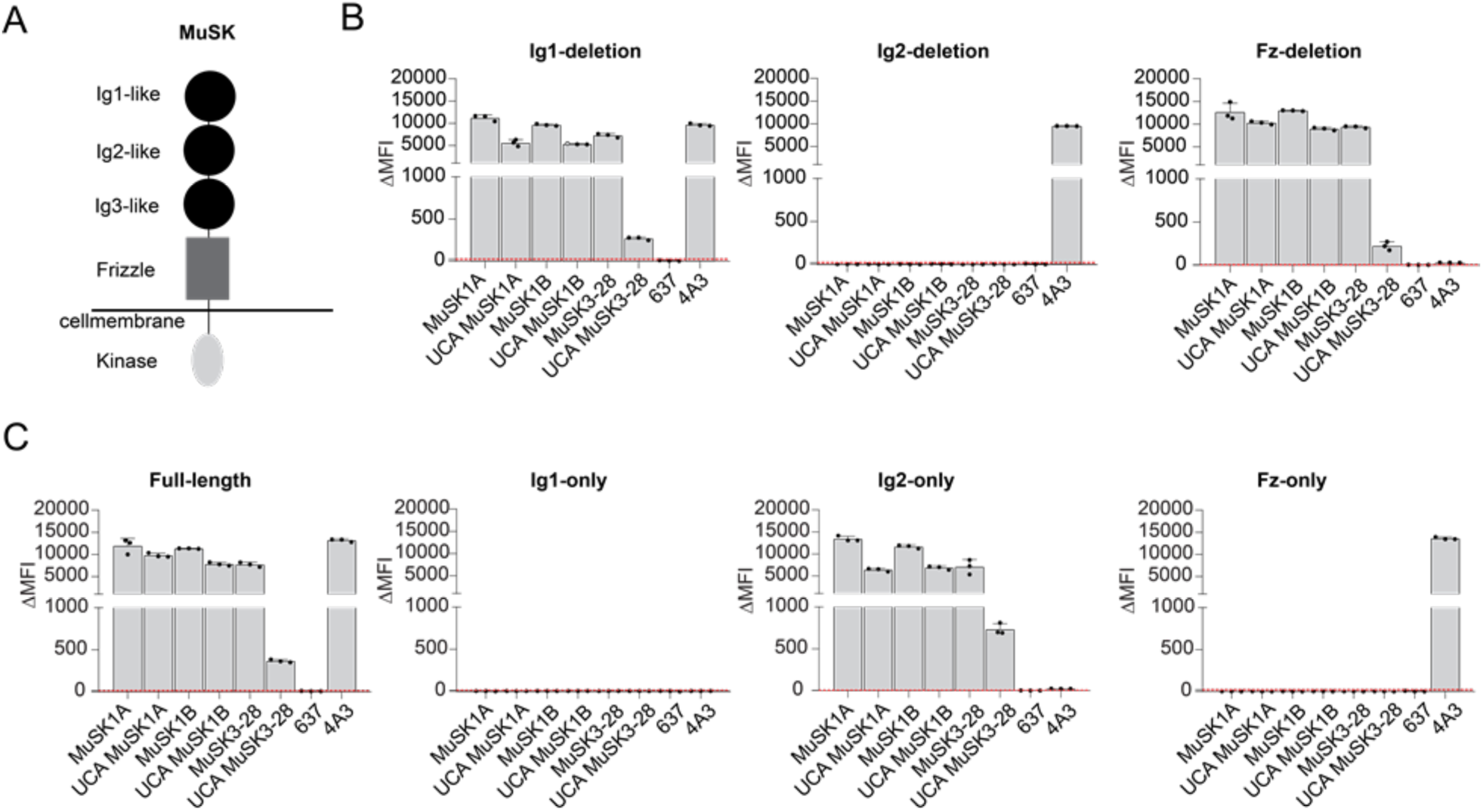
Mature MuSK mAbs and their UCA counterparts bind to the same MuSK domain. The MuSK mAbs and the UCA were tested for domain binding and recognition with a CBA expressing MuSK-GFP domain variants. **(A)** Illustration of the full-length MuSK receptor. **(B and C)** The ectodomain of MuSK consists of several different Ig-like domains and a frizzle domain. Different mutations of the MuSK protein either consisting of a domain deletion or specific domain-only construct were tested for binding by the mAbs. Humanized MuSK mAb 4A3 was used as the positive control and AChR-specific mAb 637 as the negative control. Results for each mAb are shown. The ΔMFI was calculated by subtracting the signal from non-transfected cells from the signal of transfected cells. Each data point represents a separate replicate within the same experiment, which was measured in triplicate. Bars represent means and error bars SDs. Values greater than the mean + 4SD of the negative mAb 637 indicated by horizontal dotted lines were considered positive.

**Supplemental Figure 5.**
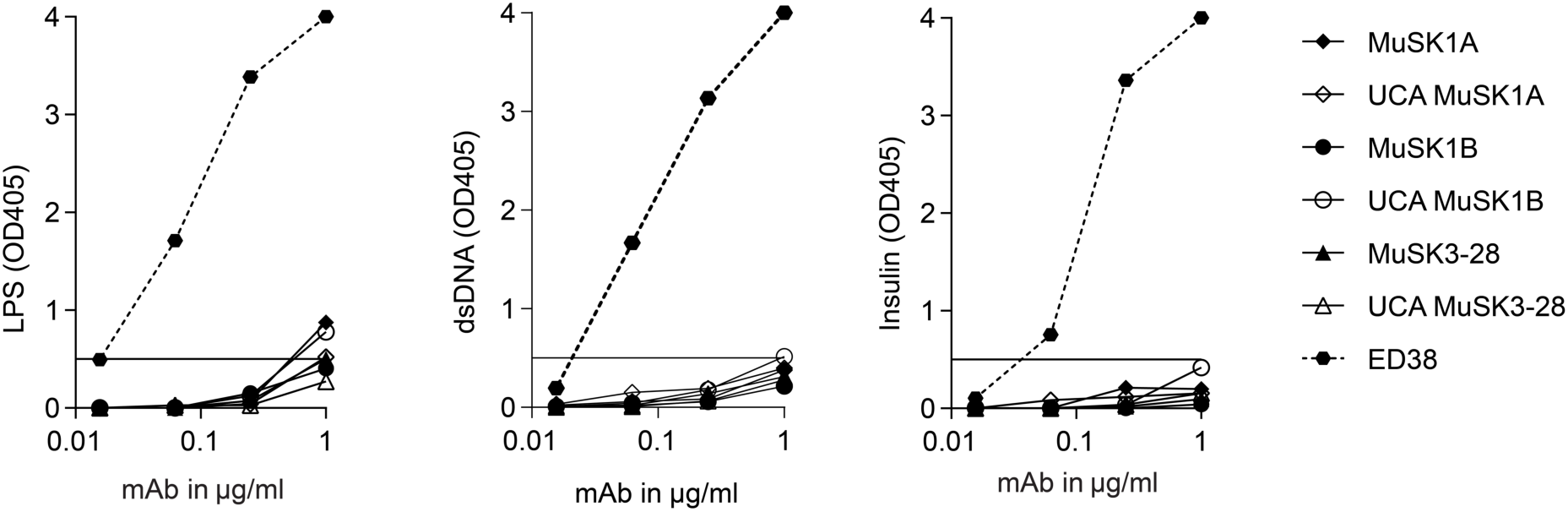
Mature and unmutated common ancestor MuSK mAbs are not polyreactive. The reactivity of the mature and UCA mAbs against LPS **(A)**, dsDNA **(B)** and Insulin **(C)** was tested by ELISA. ED38, a monoclonal antibody cloned from a VpreB + L + peripheral B cell, was used as a positive control and shown by the dotted line curves. Solid line curves represent MuSK mAbs and the UCAs. Dotted horizontal lines mark the positive reactivity cut-off at OD405 0.5.

**Supplemental Figure 6.**
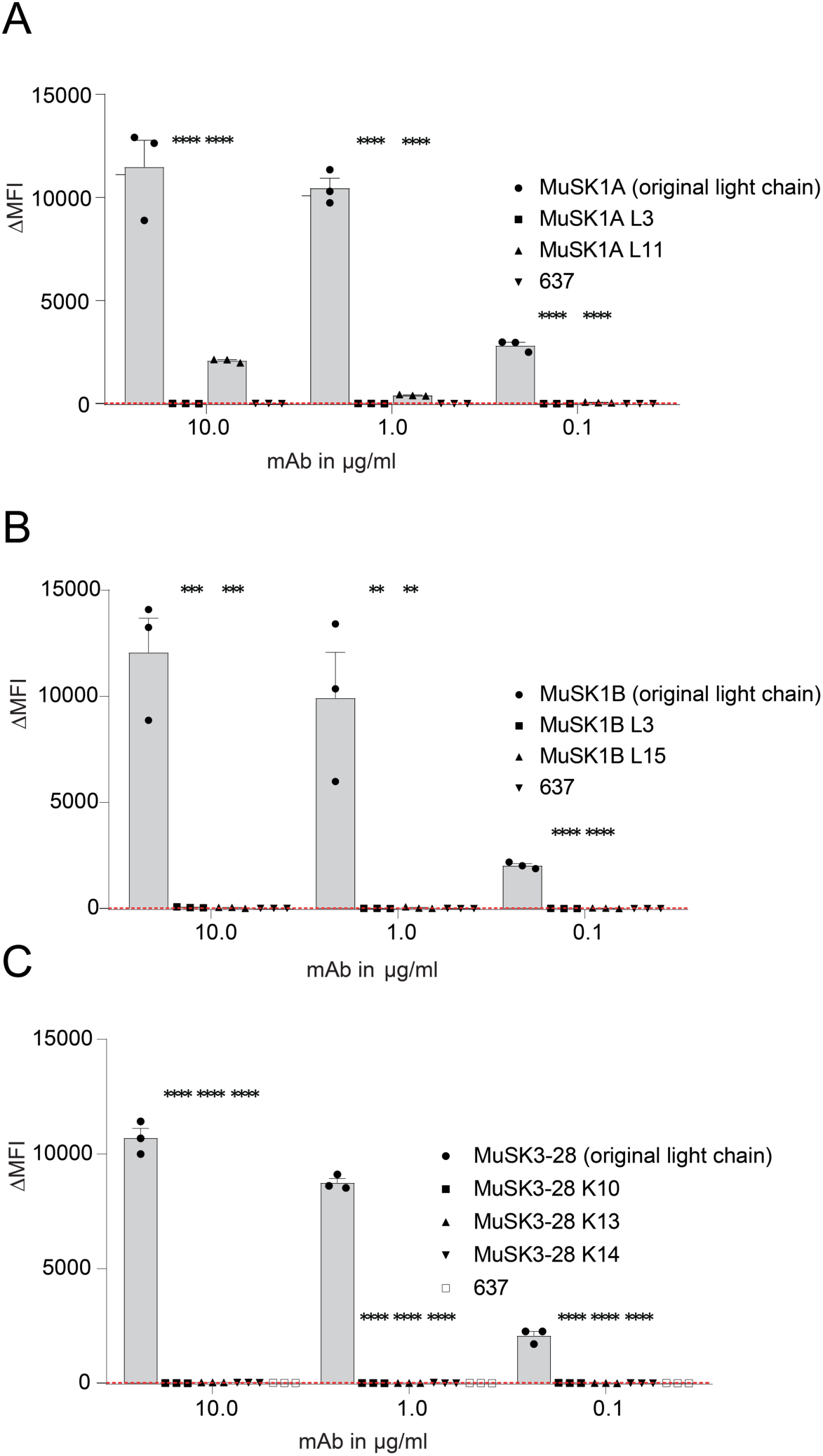
Light chains of MuSK mAbs make contributions to binding. The VL of the endogenous VH-VL pair was replaced to test the VL contributions to binding. A-C. All three mature mAbs MuSK1A **(A)**, MUSK1B **(B)** and MuSK3-28 **(C)** were paired with light chains from different subtypes consistent with their subclass and recombinantly expressed. These newly generated mAbs were tested for their binding capacity by CBA. AChR-specific mAb 637 was used as the negative control. Bars represent means and error bars SDs. The ΔMFI was calculated by subtracting the binding of the non-transfected cells from that of the transfected cells. Values greater than the mean + 4SD of the negative mAbs at 1.25 µg/ml (indicated by the horizontal dotted line) were considered positive. Statistical differences are shown when significant (Multiple comparisons ANOVA against the pooled results of the endogenous heavy and light chain combination with Dunnet’s correction; * p<0.05, ** p<0.01, *** p<0.001, **** p<0.0001).

**Supplemental Figure 7.**
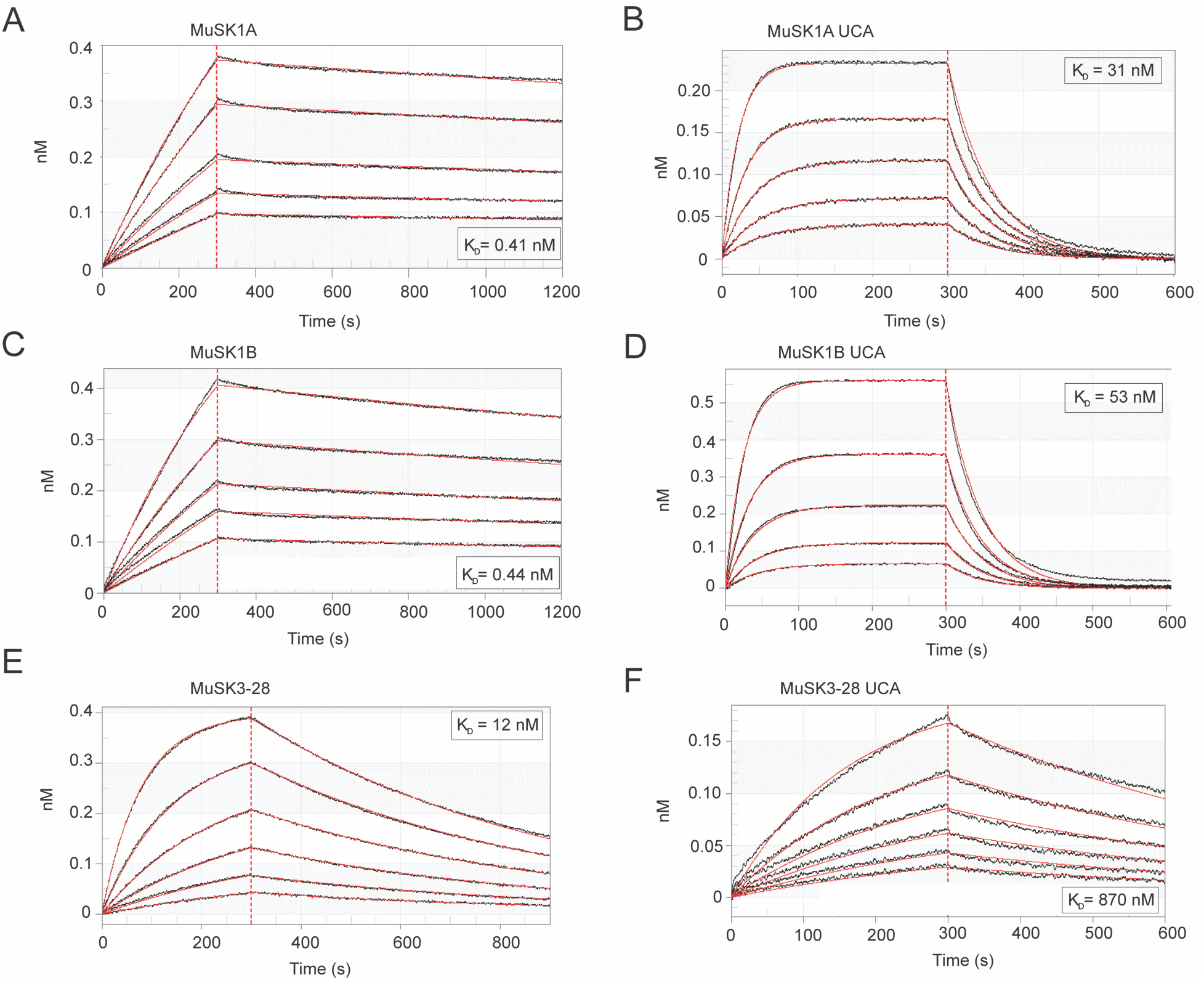
The affinity of the MuSK mAb UCAs is lower than that of the mature counterpart. Affinity of the mature and UCA Fabs to MuSK was determined by biolayer interferometry. A serial dilution series of the Fabs (900 – 1 nM) were used to determine the binding affinity with the captured MuSK. **(A – F)** Affinity measurements of the mature antibodies **(A, C, E)** and their UCA counterparts **(D, E, F)**. The X-axis depicts the time in seconds. The Y-axis depicts the wavelength shift detected by biolayer interferometry, which is proportional to material bound (nM). The K_D_ values are shown for each measurement.

**Supplemental Figure 8.**
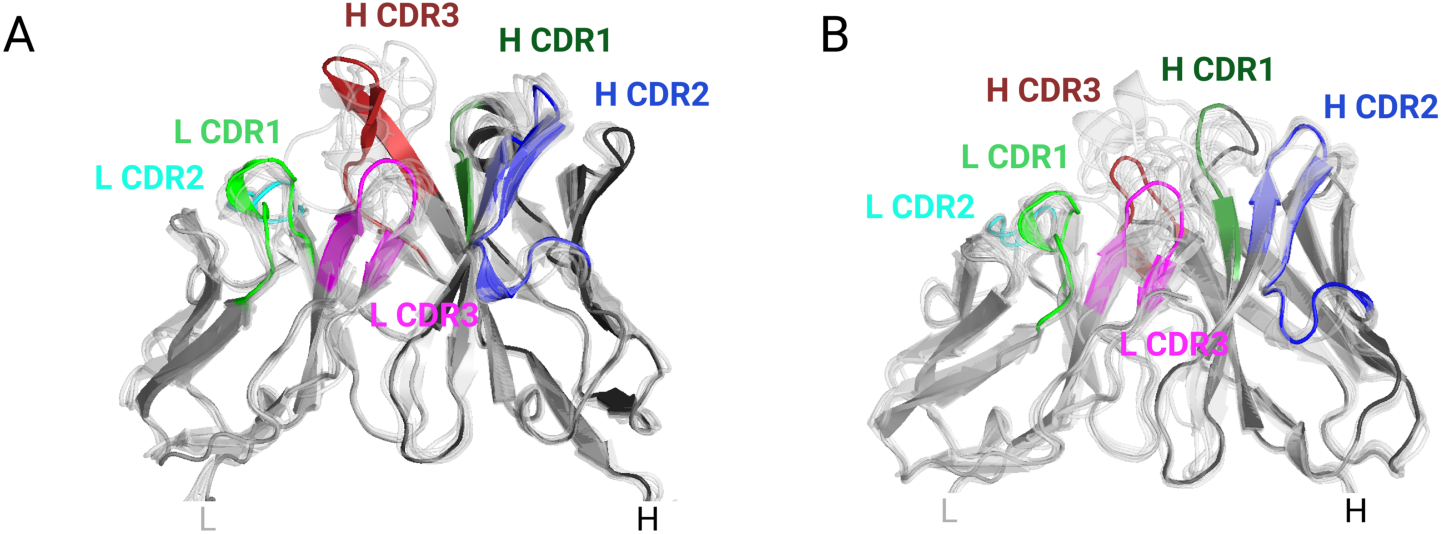
MuSK1A and MuSK1B are similar to sequence related Fabs. MuSK1A and MuSK1B differ most substantially in the L CDR3 and H CDR3 loops compared to sequence-related Fabs from the PDB. **(A)** Superposition of VH and VL domains of MuSK1A with related Fabs. PDB codes 3X3F, 3X3G, 5IES, 6DW2, 6ID4, 6MFP, 6P8N, 6MFJ and 6PHF are aligned to the VH domain. PDB codes 4DAG, 4QHK, 4QHL, 4XNM, 4XNQ, 5ODB, 5Y2K, 6B0S and 6EIB are aligned to the VL domain of MuSK1A. **(B)** Superposition of VH and VL domains of MuSK1B with related Fabs. PDB codes 1QLR, 3B2U, 4R4B, 5DRW, 5SX4, 5DRX, 6MLK, 6MHR, 6II9 and 6B3S are aligned to the VH domain of MuSK1B. PDB codes 4AJ0, 4AIX, 4DAG,4QHK, 4QHL, 4XWG, 5BV7, 5ODP, 5Y2K, 6BOS and 6EIB are aligned to the VL domain of MuSK1B.

